# Intestinal Dysbiosis Alters Acute Seizure Burden and Antiseizure Medicine Activity in the Theiler’s Virus Model of Encephalitis

**DOI:** 10.1101/2024.11.29.625527

**Authors:** Inga Erickson, Stephanie Davidson, Hanna Choi, Seongheon Rho, Michelle Guignet, Kristen Peagler, Kenneth Thummel, Aaron Ericsson, Melissa Barker-Haliski

## Abstract

**Objective:** Brain infection with Theiler’s virus (TMEV) in C57BL/6J mice produces an etiologically relevant model of acquired seizures. Dietary changes can modify acute seizure presentation following TMEV brain infection and influence intestinal microbiome diversity and composition. Intestinal dysbiosis may thus similarly affect seizure burden and antiseizure medicine (ASM) activity in this model, independent of pharmacokinetic effects. We thus sought to define the influence of antibiotic (ABX)-induced gut dysbiosis on acute seizure presentation, anticonvulsant activity of carbamazepine (CBZ), and CBZ pharmacokinetics with TMEV infection.

**Methods:** Male C57BL/6J mice (4-5 weeks) received oral (p.o.) ABX or saline (SAL) once daily beginning on arrival through Day 7 post-TMEV infection (p.i.). Mice were infected with TMEV or PBS on Day 0. Mice received intraperitoneal (i.p.; 20 mg/kg) CBZ or vehicle (VEH) twice daily Days 3-7 p.i. and were assessed for handling-induced seizures 30 min after treatment. Plasma was collected on Day 7 p.i. at 15 and 60 min post-CBZ administration for bioanalysis.

**Results:** TMEV infection induced acute seizures, but ABX-induced gut dysbiosis altered seizure presentation. There were 75% SAL-VEH, 35% SAL-CBZ, 35% ABX-VEH, and 72% ABX- CBZ mice with seizures during the 7-day monitoring period. There was a significant pretreatment x ASM interaction (p=0.0001), with differences in seizure burden in SAL- versus ABX-pretreated mice (p=0.004). CBZ significantly increased latency to seizure presentation; an effect absent in ABX-CBZ mice. Plasma CBZ concentrations did not differ between SAL and ABX pretreatment groups, suggesting that ABX did not influence CBZ pharmacokinetics.

**Significance:** ABX-induced gut dysbiosis markedly altered acute disease trajectory with TMEV-induced encephalitis, reflecting a novel contribution of the gut microbiome to seizure presentation. ABX-induced gut dysbiosis also significantly changed acute seizure control by CBZ, but did not influence plasma CBZ concentrations. The gut-brain axis is thus an under- recognized contributor to TMEV infection-induced seizures, ASM activity, and disease burden.

**Key Points:**

- Theiler’s virus infection in mice models encephalitis that exhibits differential disease trajectory with gut microbiome modulation.
- Experimentally evoked gut dysbiosis, i.e. a disrupted gut microbiome, dramatically shifts the anticonvulsant activity of carbamazepine.
- There is no concomitant shift in circulating carbamazepine concentrations in TMEV-infected mice with and without a dysbiotic microbiome.
- The gut microbiome is an under-recognized driver of seizure risk and antiseizure medicine activity in the Theiler’s virus mouse model.

## Introduction

Epilepsy is a long-term complication of viral infection-induced brain encephalitis. Patients with viral encephalitis and seizures are at higher risk of developing epilepsy post-infection (up to 22 times)^1^. Viral infection-induced epilepsy accounts for a significant portion of the total worldwide epilepsy burden, especially in low- and middle-income countries, such as Côte d’Ivoire (13%) and Mali (47%)^2^. Therefore, greater preclinical integration of viral infection-induced epilepsy models may more comprehensively and equitably address the global clinical acquired epilepsy burden.

Few animal models of infection-induced acute seizures and epilepsy exist. However, one particularly useful rodent model is evoked by brain infection with the Theiler’s murine encephalomyelitis virus (TMEV). TMEV is a non-enveloped, positive-stranded RNA virus in the *Picornaviridiae* family^3,4^. When C57BL6/J mice are infected with TMEV delivered into the brain, they develop acute, symptomatic seizures around day 3 post infection (p.i.), before clearing the virus by around day 7 p.i.^5^. A majority (∼50-65%) of mice that initially presented with acute symptomatic seizures will weeks later go on to later develop spontaneous recurrent seizures (SRS), i.e. epilepsy^6^. Further, TMEV-infected mice that initially present with acute symptomatic seizures during the infection period demonstrate reduced seizure threshold long-term and exhibit cognitive and behavioral deficits, consistent with clinical comorbidities of epilepsy^5,7,8^. Finally, mice that develop acute symptomatic seizures and epilepsy also present with extensive hippocampal damage and neuroinflammation, further revealing that the TMEV model provides an important mouse platform to investigate novel ictogenic and epileptogenic mechanisms.

The gut microbiome is increasingly recognized as an important modifier of seizure presentation, epilepsy pathology, and pharmacological treatment response^9^. The gut microbiome modulates carbohydrate and amino acid metabolism, microglial and astrocytic function, and hippocampal neurotransmitter levels^9^, all of which directly influence the progression and severity of epilepsy. Importantly, patients with drug-resistant epilepsy (DRE) have an altered gut microbiome compared to patients with well-controlled epilepsy and healthy controls^10^. Microbiome changes, i.e. dysbiosis, can also influence the behavioral response of rodent epilepsy models. For example, Medel-Matus et. al used chronic stress as a microbiome modulator in an amygdala-kindled rat model of epilepsy^11^, showing that stress-induced gut dysbiosis accelerates epileptogenesis and increases seizure severity. Indirect modulation of the gut microbiome may thus meaningfully affect epileptogenesis. Indeed, the ketogenic diet has been successfully used for over a century to treat a variety of epilepsy syndromes in humans, including reducing seizure burden and improving cognitive and motor functions in children with DRE^12^. The ketogenic diet alters the gut microbiome, and these alterations may be responsible for its anti-seizure effects^13^. Thus, environmental factors that modify the composition and diversity of the gut microbiome may indirectly influence seizure susceptibility and epilepsy burden.

Although there is high translational validity to the worldwide causes of epilepsy, intrinsic variability in disease burden with the TMEV model of infection-induced acute seizures may limit its widespread integration in preclinical research. Viral strain dramatically alters disease course and acute seizure susceptibility in C57BL/6J mice^3^. Diet can also modify the presentation of acute seizures in TMEV-infected mice^14,15^, with these dietary changes affecting long-term behavioral comorbidities of disease through potential effects on the composition of the gut microbiome^14^. While this variability may provide insight into the processes of ictogenesis and epileptogenesis, it also illustrates a major source of lab-to-lab variability that may hamper the successful integration of this model into investigational drug studies for epilepsy prevention. For example, some preliminary evidence exists to suggest that the efficacy of antiseizure medicines (ASMs) against TMEV-induced symptomatic seizures may vary between laboratories, possibly due to environmental factors. Specifically, Barker-Haliski et al. first reported that oral administration of levetiracetam (LEV; 50 mg/kg) during the active TMEV infection period was proconvulsant^16^, with LEV-treated mice having both increased cumulative seizure burden and total seizure burden. Conversely Metcalf et al. later demonstrated that a similar intraperitoneal (i.p.) dose of LEV had an anticonvulsant effect^17^, with 50% of TMEV-infected mice protected from handling-induced seizures. These two studies used a common animal source, virus source, and investigator training/protocol. The only differentiating factor was diet and environmental conditions. Subsequent studies have revealed that dietary manipulation alone can markedly change acute disease presentation^14^, suggesting that environmental influences on the gut microbiome may indirectly influence this perceived difference in LEV’s anticonvulsant activity with TMEV infection-induced acute seizures. Therefore, we hypothesized that alterations in the composition and diversity of the gut microbiome may influence acute symptomatic seizure presentation and ASM efficacy in the TMEV model without corresponding changes in ASM bioavailability and pharmacokinetics.

To fill this knowledge gap, we thus sought to define the extent to which antibiotic (ABX) administration-induced gut microbiome dysbiosis would influence the TMEV model phenotype.

We aimed to rigorously define how experimentally evoked gut dysbiosis could change acute seizure presentation, seizure-induced neuropathology, and clinical disease progression up to 7 days p.i. Secondarily, we sought to characterize the anticonvulsant activity of a commonly prescribed ASM, carbamazepine (CBZ), and the pharmacokinetic profile of this ASM in the TMEV infection-induced acute seizure model in mice to clearly demonstrate whether ASM activity is influenced by gut dysbiosis in this etiologically relevant mouse model of acute encephalitis and epileptogenesis. We herein clearly establish the contribution of the gut microbiome in shaping acute disease progression and ASM activity after brain infection with TMEV in mice. TMEV model replicates human acquired epilepsy with a high degree of fidelity. As a result, this study provides critical insight to definitively demonstrate that the gut microbiome is a novel contributor to seizure susceptibility, epileptogenesis, and ASM anticonvulsant activity. This study highlights that the gut microbiome plays an under-recognized and outsized role in shaping an individual’s risk to an epileptogenic insult and that gut dysbiosis can dramatically alter pharmacological activity of ASMs without affecting circulating plasma concentrations, all of which warrants greater mechanistic investigation.

## Materials & Methods

See Figure 1 for general experimental overview and *Supplemental File* for detailed study methods.

**Figure 1.**
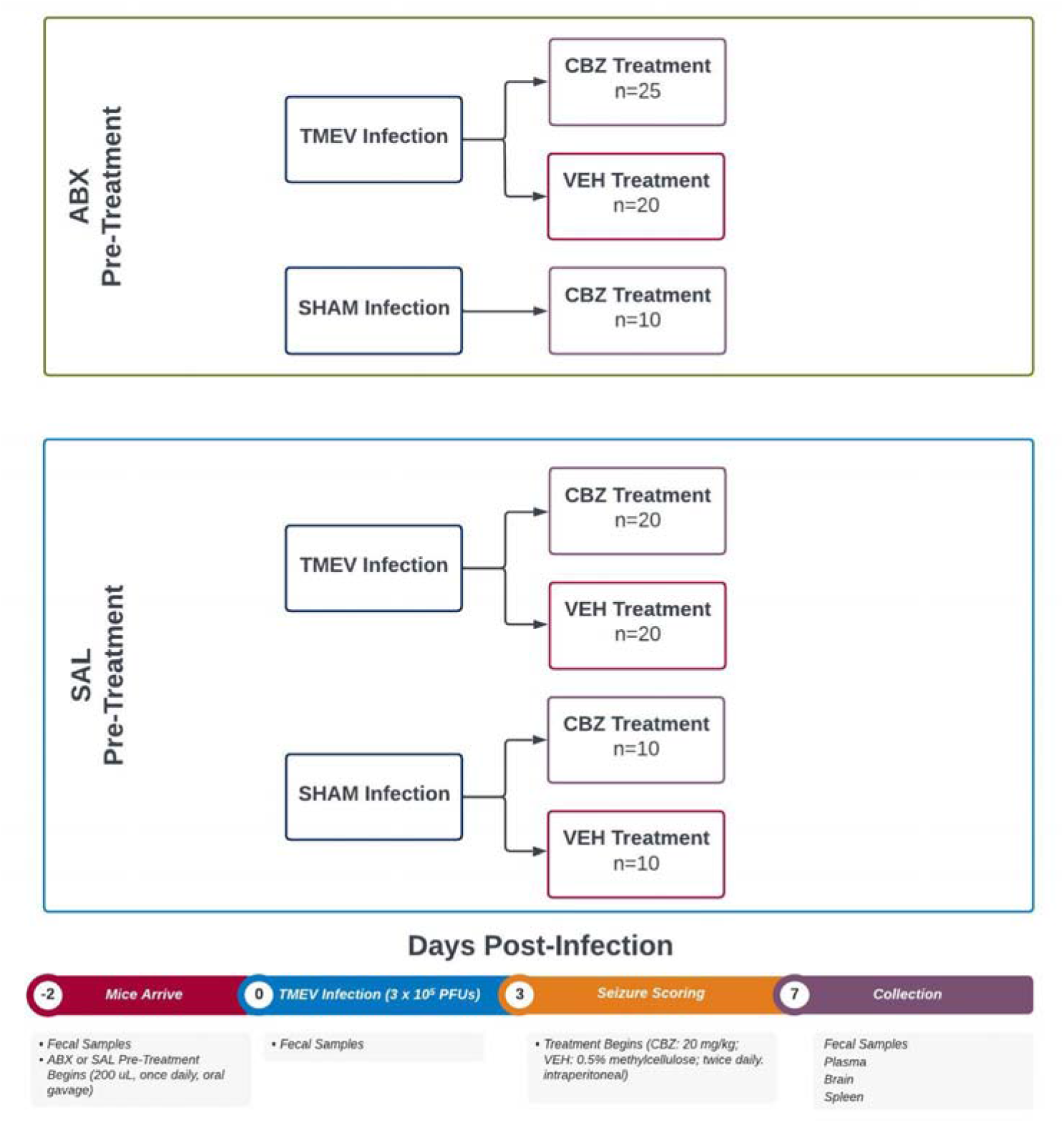
Overview of experimental design and study timeline to test the hypothesis that experimentally induced alterations in the gut microbiome using antibiotics can evoke significant differences in acute behavioral seizure presentation and anticonvulsant efficacy in the TMEV model evoked in male C57BL/6J mice (aged 4-5 weeks). The antibiotic cocktail (ABX) consisted of ampicillin (1 g/L), metronidazole (1 g/L), neomycin sulfate (1 g/L), and vancomycin (0.5 g/L) administered 1x/day by oral gavage; half of mice were randomized to saline pretreatment. The dose of carbamazepine (CBZ) was selected based on its known anticonvulsant activity in a variety of rodent seizure models, which was administered to mice twice per day 30 min prior to behavioral seizure scoring. Mice were randomized to receive either 0.5% methylcellulose vehicle (VEH) or CBZ from day 3 post-infection within the brain with Theiler’s murine encephalomyelitis virus (TMEV). On the final testing day, animals were euthanized at discrete time points after CBZ or VEH administration to assess potential that ABX-induced gut microbiome dysbiosis could influence pharmacokinetics of CBZ.

## Results

### Intestinal dysbiosis disrupts TMEV-induced seizure presentation

Intestinal dysbiosis in the mouse TMEV model was experimentally induced because this seizure model is sensitive to dietary modification, potentially due to changes in the gut microbiome^14^. Sham-infected mice maintained or gained weight, whereas TMEV infection evoked acute body weight loss (Figure 2A; time x treatment group interaction - F (54, 971) = 4.84, p<0.0001). Relative to SAL-VEH sham-infected mice, ABX administration exacerbated body weight loss through 4 days p.i., but this weight loss continued in TMEV-infected mice through 10 p.i. (Figure 2A; p<0.05 for all ABX-pretreated mice). ABX-pretreated sham infected mice regained body weight by 5 days p.i. such that there was no difference from SAL-VEH sham-infected mice. Conversely, SAL-pretreated mice infected with TMEV began to experience significant body weight loss beginning on 4 days p.i. through the study end (Figure 2A; p<0.05 for all SAL-pretreated mice). No other groups demonstrated significant differences from SAL-VEH sham-infected mice during the 10-day monitoring period.

**Figure 2.**
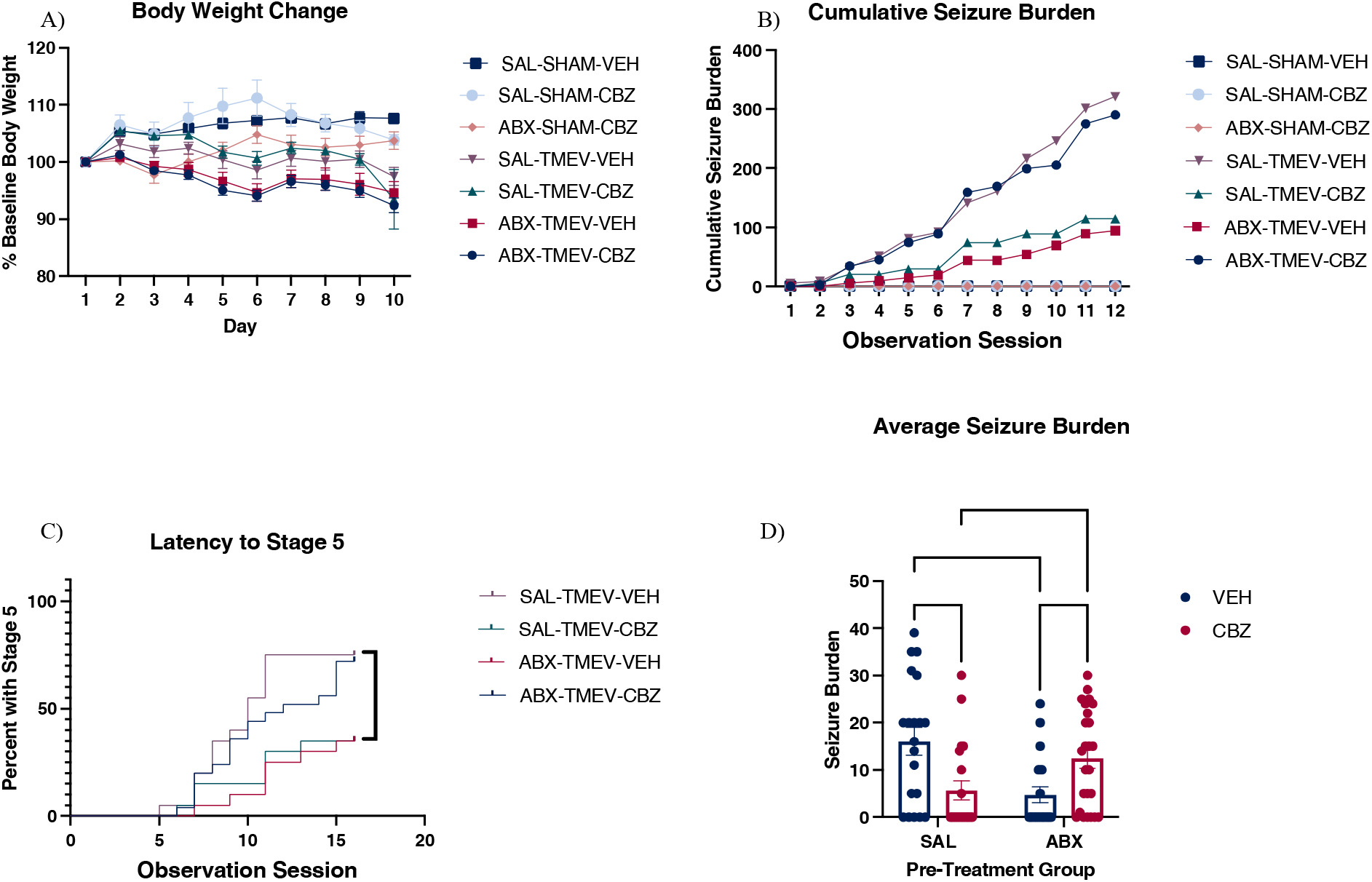
Intestinal dysbiosis with a 10-day course of oral antibiotics cocktail (ABX) disrupts TMEV-induced acute symptomatic seizure presentation from days 3 to 7 post-infection in male C57BL/6J mice aged 4-5 weeks old. Mice were monitored for behavioral seizures twice per day. A) TMEV infected mice lose weight during the acute infection period. B, C) In mice with an intact microbiome, carbamazepine (CBZ) significantly decreases seizure burden compared to VEH. In mice with ABX-induced gut dysbiosis, CBZ significantly increases seizure burden compared to VEH. D) In mice with an intact gut microbiome, CBZ increases the latency to a Stage 5 seizure; in mice with ABX-induced gut dysbiosis, CBZ decreases the latency to a Stage 5 seizure. Data analyzed with Mantel-Cox Log rank test (C) and two-factor ANOVA (D). * = P = 0.03, ** = P = 0.002, *** = P = 0.0002.

Sham-infected mice did not have any acute symptomatic seizures (Figure 2B), but TMEV infection induced acute symptomatic seizures, regardless of ABX pretreatment and ASM history. Comparing specific treatments during the twice daily seizure monitoring sessions, in mice with an intact gut microbiome, VEH-treated mice had more seizures versus CBZ-treated mice throughout the study monitoring period (Figure 2B). In TMEV-infected mice with ABX-induced intestinal dysbiosis, VEH-treated mice had a lower cumulative seizure burden relative to SAL-VEH mice. ABX-mediated intestinal dysbiosis was associated with reduced seizure burden. CBZ-treated mice with intestinal dysbiosis had a significantly higher cumulative seizure burden.

Further, the latency to the first observed Racine stage 5 seizure was substantially increased by repeated CBZ administration during days 3-7 p.i.; an effect that was not present in ABX-treated mice similarly treated with CBZ (Figure 2C). In mice with an intact microbiome, CBZ reduced the number of mice that developed a stage 5 seizure whereas this effect was reversed in mice with gut dysbiosis. Further in TMEV-infected mice with intestinal dysbiosis, VEH treatment was associated with fewer observed seizures compared with CBZ treatment (Figure 2D). There was a significant pretreatment x ASM interaction effect (F (1, 81) = 16.0, p=0.0001), with post-hoc tests revealing marked differences in seizure burden in SAL- versus ABX-pretreated mice (p=0.0001). Conversely, the average seizure burden of mice treated with CBZ with an intact microbiome was significantly lower than the average seizure burden of mice with an intact microbiome treated with VEH. This demonstrates that CBZ has an acute anticonvulsant effect, consistent with prior reports^7^. In mice with intestinal dysbiosis, the average seizure burden was significantly higher with CBZ treatment than with VEH treatment, demonstrating that in the setting of gut dysbiosis in the TMEV model, CBZ instead evokes a proconvulsant effect. Altogether, experimentally evoked gut dysbiosis disrupted TMEV infection-induced acute seizure presentation and acute seizure control by CBZ.

### Markers of overall health were not impacted by antibiotic administration

Blood-based markers of overall health were assessed via a routine veterinary complete blood count panel to rule out potential peripheral pathology or unanticipated effects of TMEV infection and/or repeated ABX administration. Complete blood counts revealed that analytes were similar across saline and ABX pretreatment groups (Figure 3) within both sham- and TMEV-infected groups, suggesting that a 10-day course of oral ABX administration did not adversely affect any leukocyte or erythrocyte parameters. However, CBZ administration was associated with a significant increase in the percent of blood cells that were neutrophils, regardless of TMEV-infection history (Figure 3C; F (3,20) = 19.2, p<0.0001).

**Figure 3.**
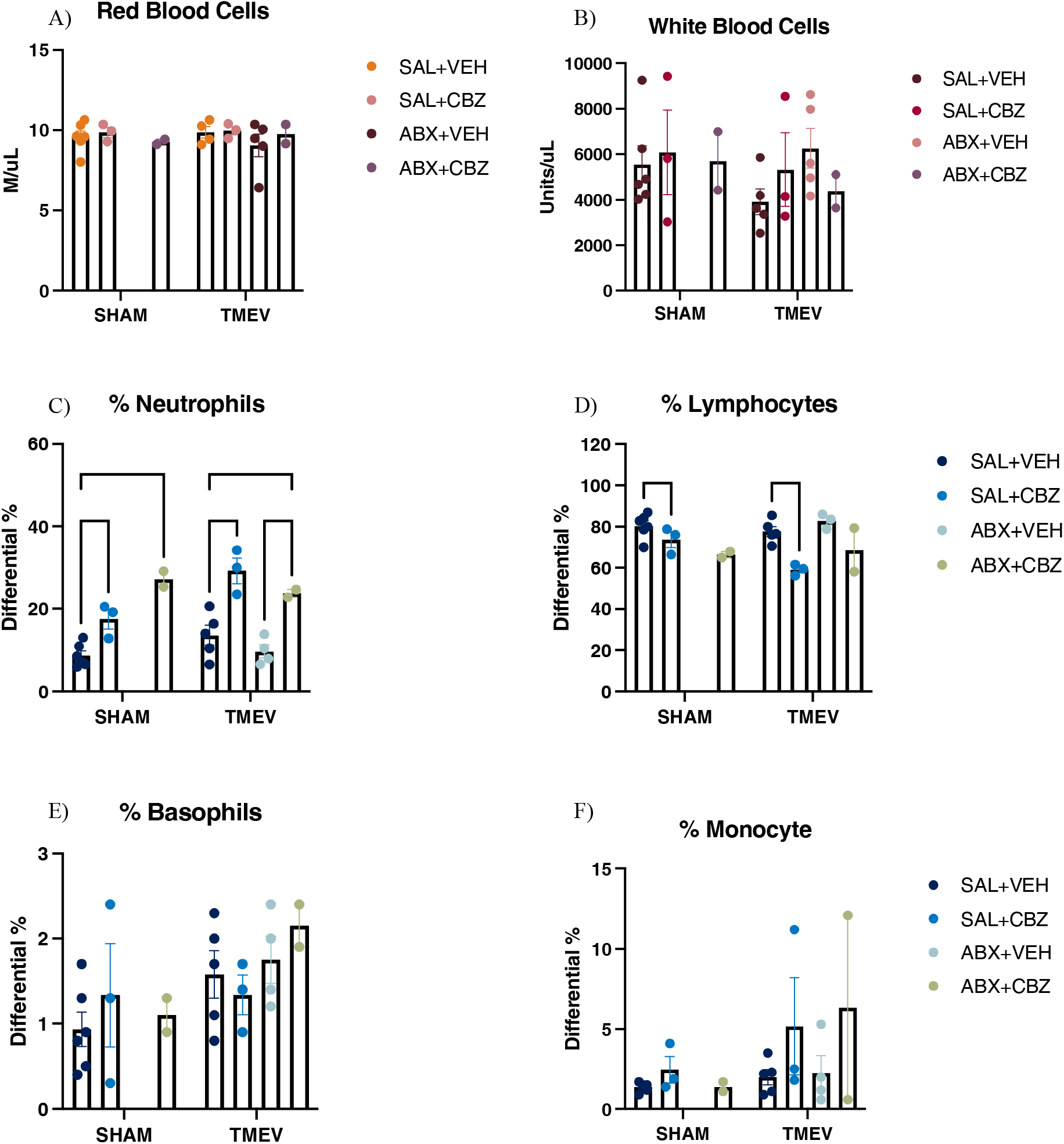
A 10-day course of oral antibiotic (ABX) administration to male C57BL/6J mice aged 4-5 weeks old infected with or without TMEV did not impact peripheral markers of overall health using a routine veterinary complete blood counts analysis. Using a veterinary diagnostic panel of complete blood count (A, B) and differential analyses (C-F) indicate that mice are clinically normal, other than brain TMEV infection. There is an effect of repeated CBZ administration on the differential levels of lymphocytes (D) and basophils (E), but no effect of repeated ABX administration on any outcome measure. Data analyzed with Tukey’s multiple comparisons test. * = P = 0.03, *** = P = 0.0002.

While clinical pathology biomarkers were not adversely impacted by TMEV infection or repeated ABX administration, we also wanted to confirm that immune system function was not grossly affected by ABX-induced gut microbiome depletion. TMEV infection generally acutely reduced spleen weight (F(1,81) = 29.6, p<0.0001), with a trend for an effect of repeated CBZ administration (p=0.06; Figure 4). Except for SAL-SHAM-CBZ mice, spleen weights were within the standard range for C57BL6/J mice^18^. These findings altogether indicate that mice were otherwise clinically normal, despite brain infection, and that repeated ABX administration to induce gut dysbiosis did not significantly impact clinical signs of chronic disease.

**Figure 4.**
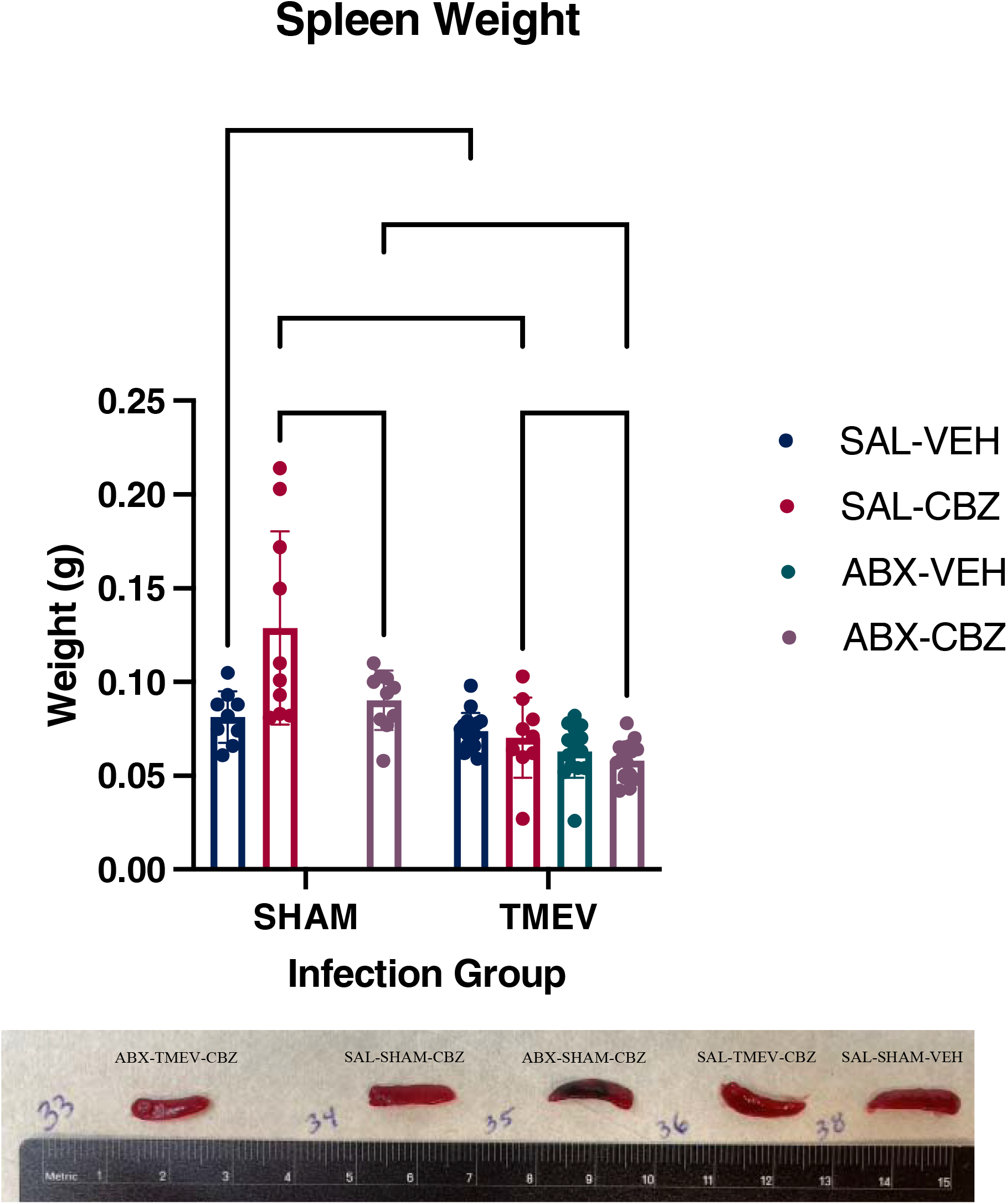
TMEV infection and carbamazepine (CBZ) treatment are associated with reduced spleen weight in male C57BL/6J mice aged 4-5 weeks old with and without a history of 10-day antibiotics (ABX) administration to deplete the gut microbiome. Data analyzed with Sidak’s post hoc test. ** = P = 0.002, **** = P = <0.0001.

### A 10-day course of oral antibiotics significantly alters the makeup of the gut microbiome

The composition and diversity of the gut microbiome was quantified at several points in the TMEV infection period to ensure that repeated ABX administration induced gut dysbiosis. Differential abundance testing was performed using serial two-factor ANOVA (Table S1), revealing multiple families that were affected by both time and treatment (interaction effects). There were several significant differences in the microbial composition between SAL pre-treated and ABX pre-treated mice across a diversity of Gram-positive and Gram-negative bacterial families, as well as differences in ABX pre-treated animals from Day −2 to Day 7 (Figure 5A).

**Figure 5.**
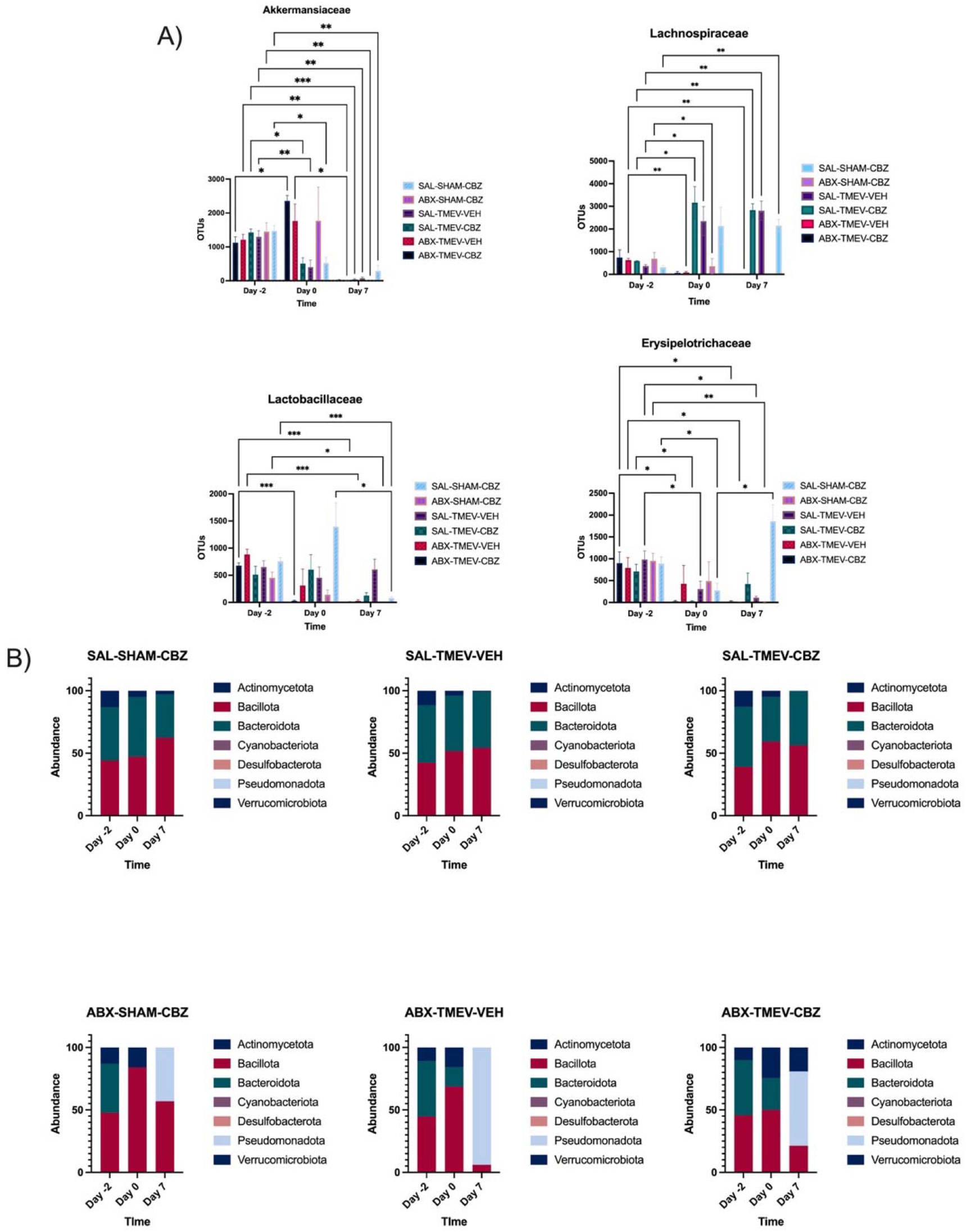
A 10-day course of oral antibiotics (ABX) administered to TMEV-infected or sham-infected male C57BL/6J mice aged 4-5 weeks old significantly altered the composition of the gut microbiome with or without repeated twice-daily carbamazepine (CBZ) administration prior to each behavioral seizure observation session. There were significant differences at the family (A) and phylum (B) levels, as well as across time. OTUs = operational taxonomic units. * = P = 0.03, ** = P = 0.002, *** = P = 0.0002, **** = P = <0.0001.

Specifically, we found that the variation in several key species known to be affected by epilepsy or other CNS disease states were particularly modified in a time x treatment group interaction effect, including *Akkermansiaceae*, *Lachnospiraceae*, *Lactobacillaceae*, and *Erysipelotrichaceae* (Figure 5A and Table S1). Further, proportional distribution of all species in each treatment group exhibited marked shifts over time (Figure 5B); the gut microbiome of SAL pre-treated animals was predominantly made up of Bacillota (Firmicutes)^19^ and Bacteroidota, across all testing days (Figure 5B). The gut microbiome of ABX pre-treated animals shared this makeup on Days −2 and 0, but on Day 7 there was a significant increase in Pseudomonadota (Proteobacteria) (Figure 5B). This demonstrates that the ABX cocktail successfully induced intestinal dysbiosis.

### Acute TMEV infection leads to hippocampal cell death

Hippocampal histopathology was performed to assess the degree of TMEV infection-induced neurodegeneration using FJ-C staining (Figure S1 and Table S1) and to determine whether ABX-mediated gut dysbiosis led to any qualitative changes in neuropathology. Sham-infected mice did not experience hippocampal cell death. Conversely, TMEV-infected mice that had seizures experienced significant cell death in hippocampal CA1 and CA3 regions, as well as overlaying somatosensory cortex. No mice experienced cell death in the dentate gyrus region of dorsal hippocampus (Figure S1 and Table S2). Of note, however, was the observation that there were fewer SAL-TMEV mice treated with CBZ with appreciable cell death versus SAL-TMEV- VEH and ABX-TMEV-VEH in area CA1 (X^2^ = 2.55, p=0.011, Table S1), likely reflective of the reduced seizure burden in this group. Altogether, these findings confirm earlier reports that the TMEV infection leads to hippocampal cell death^8^. However, there was no qualitative difference in the extent of neuropathology in ABX- or SAL-pretreated mice infected with TMEV; mice with seizures still had hippocampal cell death regardless of pretreatment history.

### Antibiotic-induced intestinal dysbiosis does not alter plasma carbamazepine concentration

Time-related plasma CBZ concentration was assessed at 7 days p.i. to determine if there was an interaction between 10-day ABX treatment and repeated CBZ administration from 3-7 days p.i. with TMEV. There was a main effect of time post CBZ administration (F (1.257, 23.26) = 9.48, p=0.0032) but there was no effect of ABX-pretreatment or time x pretreatment interaction on CBZ concentration in plasma at this time point. Thus, a 10-day course of oral ABX administration did not affect the plasma concentration of CBZ in mice (Figure 6), suggesting that the differential seizure burden observed in ABX+CBZ versus VEH+CBZ mice infected with TMEV is due to differences in the diversity and composition of the gut microbiome that affect processes largely unrelated to oral CBZ pharmacokinetic disposition (Figure 5).

**Figure 6.**
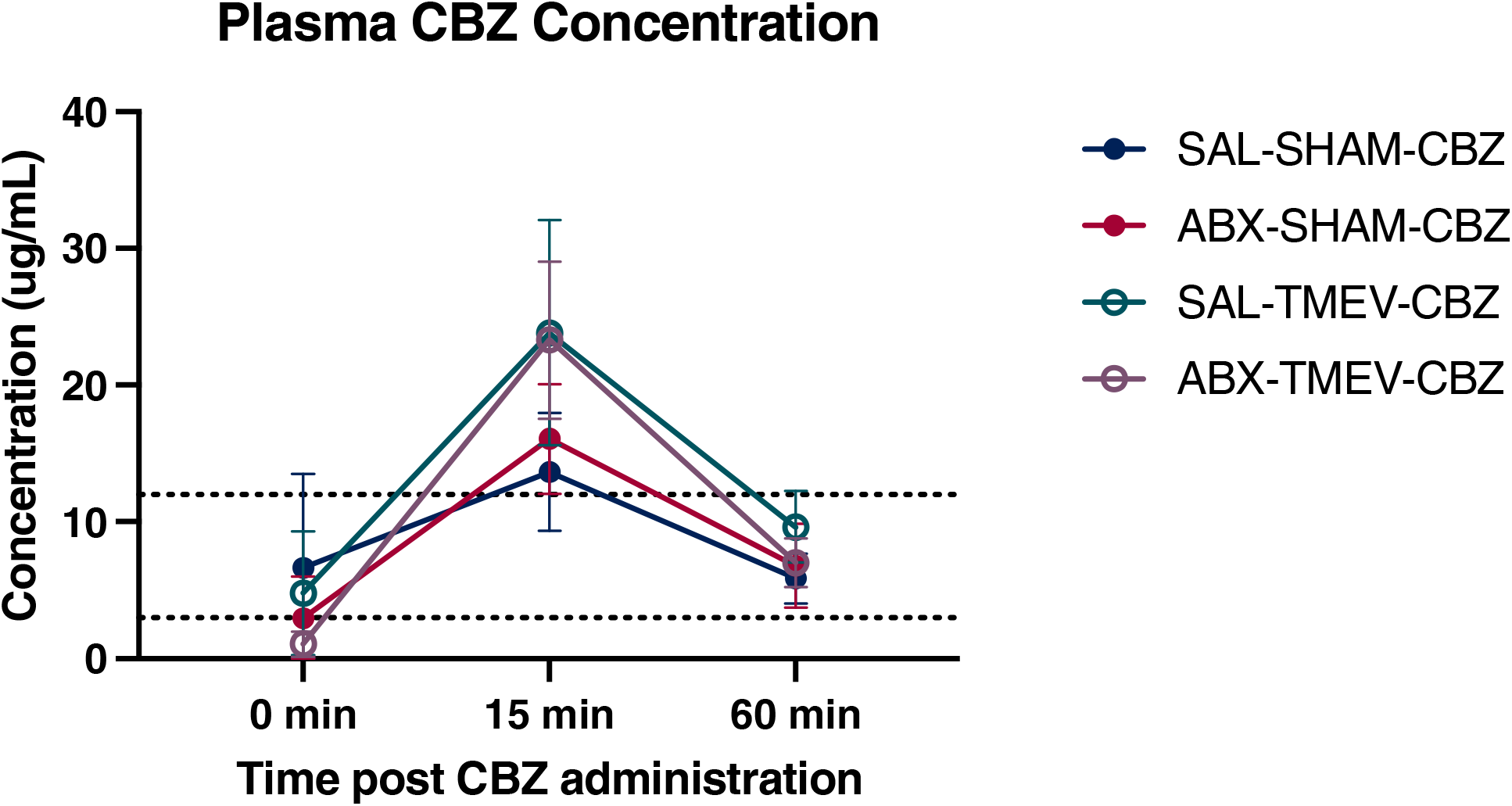
A 10-day course of oral antibiotics (ABX)-induced intestinal dysbiosis did not alter plasma carbamazepine (CBZ) concentrations (n=3-9 mice/treatment group) in male C57BL/6J mice aged 4-5 weeks old at 7 days post brain infection with TMEV to induce acute encephalitis and symptomatic seizures. Mice received 20 mg/kg (i.p.) CBZ twice per day for days 3-7 post-infection, administered 30 min prior to behavioral assessment of handling-induced seizures. On the 7^th^ day post-infection, mice were euthanized within each treatment group for timed collection of terminal blood samples for pharmacokinetic analysis by liquid chromatography/tandem mass spectrometry. Dashed lines represent the anticonvulsant range of CBZ in male CF-1 mice in other seizure tests^44^. There were no statistically significant differences in plasma concentrations across treatment groups, as measured by two-factor ANOVA.

## Discussion

This study is the first to rigorously establish that the diversity and composition of the gut microbiome is a significant source of variability in the behavioral phenotype of the TMEV model of infection-induced acute seizures. We herein now demonstrate that experimentally evoked gut dysbiosis robustly influences the severity and onset of acute symptomatic seizures in the TMEV model itself. Specifically, our study demonstrates that a 10-day course of oral ABX administration commencing prior to TMEV infection was sufficient to evoke decreased cumulative seizure burden relative to SAL pretreatment, reflective of an anticonvulsant effect of gut dysbiosis. Further, we herein demonstrate that the anticonvulsant activity of CBZ is dramatically impacted by short term gut dysbiosis in the TMEV model of acute symptomatic seizures. This study therefore reveals that the gut microbiome may play a previously unrecognized and outsized influence on the anticonvulsant activity of ASMs because the addition of CBZ administration to mice with ABX-induced dysbiosis increased the cumulative seizure burden. This trend was also evident with average seizure burden and latency to Stage 5 seizure outcome measures. Importantly, there were no clinically significant differences in markers of overall health between experimental groups, illustrating that gut dysbiosis did not change other factors related to clinical presentation. For example, histopathology as assessed at 7 days p.i. in the brains of mice with and without TMEV-infection demonstrated neuronal death consistent with previous TMEV studies. Further, hippocampal cell death was not markedly changed by ABX-induced gut dysbiosis; the biggest driver of the extent of hippocampal cell death was acute symptomatic seizure history. Despite these differences in behavioral seizure presentation across groups with and without an intact microbiome, plasma CBZ concentrations did not markedly differ between in SAL and ABX-pretreated animals. These results altogether indicate that intestinal dysbiosis is responsible for the presently observed changes in acute seizure burden and ASM efficacy.

While the ABX pretreatment regimen successfully induced gut dysbiosis (Figure 5), it did not decrease the presence of all organisms. There were increases in some Gram-positive and anaerobic bacteria^20^ in the ABX-treated mice, which may be a result of antibiotic selectivity or the extreme treatment paradigm (10-days of four different compounds). We observed significant changes in many bacterial species in ABX-treated mice. However, few of these organisms have been studied in the context of epilepsy or other neurological diseases. Some organisms that were notably altered included *Akkermansiaceae*, *Erysipelotrichaceae*, *Lachnospiraceae*, and *Lactobacillaceae*, all of which have been earlier reported to be relevant to epilepsy^13^ or other neurological disorders^21^. The ketogenic diet has been proven to be a very useful treatment strategy for DRE^12^, and an increase in intestinal *Akkermansia* and *Erysipelotrichaceae* is observed with ketogenic diet administration to mice^13^. Additionally, oral treatment with *Akkermansia* in combination with *Parabaceteroides* is able to confer seizure protection to mice fed a control diet^13^. While not yet assessed in epilepsy, reduced intestinal levels of *Lachnospiraceae* have been detected in both clinical depression^22^ and multiple sclerosis^21^, likely due to effects on the immune system^23^. Oral *Lactobacillaceae* treatment in mice can induce cortical region-dependent alterations in GABA neurotransmitter levels and may be associated with tight junction integrity and modulation of afferent sensory nerves^24,25^. Additionally, all these organisms have been implicated in butyrate production pathways^26–28^, which is a known anticonvulsant byproduct of the ketogenic diet^29^. With many studies finding relationships between these organisms and neurological function, intestinal microbes likely play a major and underappreciated role in modulating an individual’s seizure risk and epilepsy progression following an epileptogenic insult. In this regard, our present study now rigorously demonstrates that shifts in the composition of the gut microbiome can measurably shift seizure presentation in response to an acute neurological insult, further illustrating that the gut microbiome is an under- recognized, yet modifiable, contributor to seizure risk and epileptogenesis.

We selected CBZ as the ASM for this study because it has predictable and consistent clinically meaningful anticonvulsant activity in a variety of seizure and epilepsy models^16,30^. However, we have previously reported that CBZ administration can worsen seizure burden in the TMEV model despite evidence of acute anticonvulsant effects on evoked seizures shortly after drug administration^7^. CBZ alone acts as an anticonvulsant through voltage-gated sodium channel blockade in rodent models and clinical epilepsy^31^. However, our present study demonstrates that CBZ in combination with experimentally induced gut dysbiosis acts as a proconvulsant in the TMEV model of acute symptomatic seizures. Of note, CBZ administration in SAL-pretreated mice infected with TMEV was associated with reduced hippocampal cell death in SAL pre-treated mice largely because of the reduced seizure burden in this experimental group. However, there was no marked change in the numbers of CBZ-treated mice with ABX-mediated gut dysbiosis with hippocampal cell death. Indeed, acute symptomatic seizures after TMEV infection are required for neurological damage in hippocampal structures^8^. Further, we sought to demonstrate that changes in plasma CBZ concentration were not responsible for any of the observed in vivo shifts in ASM activity, finding that plasma CBZ concentrations were not different between SAL- and ABX- pretreated mice. Nonetheless, these findings altogether suggest that differences in seizure burden were not due to changes in CBZ bioavailability or metabolism because of repeated ABX administration.

The variability in anticonvulsant activity of CBZ in this study is reminiscent of the differences in LEV activity between studies by Barker-Haliski et. al and Metcalf et. al^16,17^. It is possible that difference between the husbandry condition and diets, previously shown to impact the gut microbiome in the TMEV model^14^, also had an impact on the ASM activity between these two prior LEV studies conducted in different laboratories. However, we did not presently detect a statistically significant difference in CBZ plasma concentrations in any treatment group at the 15 min post-administration time point (Figure 6), even though elevated plasma concentration under inflammation conditions have been previously observed with other ASMs, such as perampanel^32^. Nonetheless, future studies may need to assess the potential for drug-drug interactions between metronidazole and CBZ^33^ in TMEV-infected mice, especially with regard to the expression of CYP metabolic enzymes. CYP3A4 is the primary metabolizing enzyme responsible for CBZ biotransformation^34^ and inflammation itself can affect the activity of CYP enzymes, leading to clinically significant changes in a phenotypic ASM response^35^. TMEV infection is associated with increased IL-6^36^, and increased cytokine expression is known to reduce CYP3A4 activity^37,38^. Inflammation can increase the area under the curve (AUC) of CYP3A4 metabolized drugs by up to 50%, as well as increasing the Cmax and Tmax^39^. These measures indicate that drug exposure is likely higher in inflammatory conditions. CBZ is a low extraction ratio drug (0.16^40^), with high protein binding (25% unbound^41^), but repeated CBZ administration as implemented in this study would be likely also capable of inducing CYP3A4 enzyme expression. These pharmacokinetic parameters altogether indicate that the anticonvulsant activity of CBZ would be even more susceptible to the metabolic changes induced by inflammation^39^. Systemic inflammation may explain the trend towards increased plasma concentration of CBZ in TMEV-infected mice, although this difference was not statistically significant (Figure 6).

The TMEV model is one of the few models of infection-induced acquired epilepsy, and as such is used in the National Institute of Neurological Disorders and Stroke’s Epilepsy Therapy Screening Program to profile novel investigational treatments for epilepsy^17^. However, we now definitively demonstrate that gut dysbiosis markedly alters the presentation of symptomatic seizures and acute disease burden in the TMEV model (Figure 2). The variability in this model may negatively affect the perceived efficacy of investigational ASMs, and care needs to be taken to ensure that the results are interpreted in context. Using experimental ABX administration, we now reveal that changes in the gut microbiome clearly alter seizure burden in untreated and ASM-treated mice in this acute seizure model. In fact, this study further illustrates that the gut microbiome may be a relevant therapeutic target for seizure control and that the acute seizure phenotype and ASM response of the TMEV model is particularly sensitive to experimental manipulation of the intestinal microbiome.

The gut-brain axis is altogether an understudied target in epilepsy that may benefit from greater investigation to truly define the extent to which clinical management and individual epilepsy risk can be affected by the gut microbiome. There are still numerous gaps in understanding how to harness dietary and environmental factors to influence ASM activity and seizure burden. Yet, there is potential for modulation of the gut microbiome as a treatment for epilepsy. Some epilepsy patients experience seizure freedom, or a significant decrease in seizure burden, during a short course of ABX treatment^42^. There is also some evidence that probiotic supplementation can reduce seizure burden^43^. However, there is limited clinical and preclinical research on the direct and indirect contributions of the gut microbiome on seizure susceptibility and epileptogenesis. Additional studies should thus include further rigorous investigation to identify novel therapeutic strategies that can transform the gut microbiome from an interesting experimental paradigm into a transformative disease modifying treatment for people at risk of developing epilepsy.

## Data Availability

16s sequencing data is available through the NCBI Bioproject. All other clinical monitoring and behavioral data is available upon request from the corresponding author.

## Funding Acknowledgments

Plein Center for Aging and University of Washington School of Pharmacy.

