## Supplemental Information for "Intestinal Dysbiosis Alters Acute Seizure Burden and Antiseizure Medicine Activity in the Theiler’s Virus Model of Encephalitis"

**Supplemental Tables.**

**Table S1: Differential Abundance of Intestinal Microbiome.** A 10-day course of oral antibiotics (ABX) administered to TMEV-infected or sham-infected mice significantly altered the composition of the gut microbiome with or without repeated twice-daily carbamazepine (CBZ) administration prior to each behavioral seizure observation session. Results represent the two-factor ANOVA analysis for all significant time x treatment interaction effects and p values. Values reported in ***bold italics*** are further represented in Figure 5 for individual group changes

| **Phylum** | **Family** | **P value** | **F** |
| --- | --- | --- | --- |
| *Actinobacteriota* | *Bifidobacteraceae* | P=0.4570 | F (10, 48) = 1.000 |
|  | *Microbacteraceae* | P=0.4570 | F (10, 48) = 1.000 |
|  | *Propionibacteraceae* | P=0.6445 | F (10, 48) = 0.7832 |
|  | *Atopobiaceae* | P=0.4570 | F (10, 48) = 1.000 |
|  | *Coriobacteriales Incertae Sedis* | P=0.0166 | F (10, 48) = 2.500 |
|  | *Eggerthellaceae* | P=0.4794 | F (10, 48) = 0.9722 |
| *Bacteroidota* | *Muribaculaceae* | P<0.0001 | F (10, 48) = 5.671 |
|  | *Rikenellaceae* | P=0.4570 | F (10, 48) = 1.000 |
|  | *Chitinophagaceae* | P=0.4570 | F (10, 48) = 1.000 |
| *Cyanobacteria* | *Chloroplast* | P=0.4750 | F (10, 48) = 0.9776 |
| *Desulfobacterota* | *Desulfovibrionaceae* | P=0.4570 | F (10, 48) = 1.000 |
| *Firmicutes* | *Acholeplasmataceae* | P=0.0109 | F (10, 48) = 2.677 |
|  | *Bacillaceae* | P=0.0698 | F (10, 48) = 1.891 |
|  | *Planococcaceae* | P=0.0011 | F (10, 48) = 3.644 |
|  | *Erysipelatoclostridiaceae* | P=0.0011 | F (10, 48) = 3.666 |
|  | *Erysipelotrichaceae* | ***P<0.0001*** | F (10, 48) = 4.905 |
|  | *Carnobacteriaceae* | P=0.2456 | F (10, 48) = 1.323 |
|  | *Enterococcaceae* | P<0.0001 | F (10, 48) = 4.864 |
|  | *Lactobacillaceae* | ***P=0.0025*** | F (10, 48) = 3.305 |
|  | *Leuconostocaceae* | P=0.0422 | F (10, 48) = 2.107 |
|  | *Paenibacillaceae* | P=0.4570 | F (10, 48) = 1.000 |
|  | *RF39* | P=0.0025 | F (10, 48) = 3.313 |
|  | *Staphylococcaceae* | P<0.0001 | F (10, 48) = 15.16 |
|  | *Clostridia vadinBB60 group* | P=0.1212 | F (10, 48) = 1.650 |
|  | *Clostridia UCG-014* | P=0.1040 | F (10, 48) = 1.717 |
|  | *Clostridiaceae* | P=0.0375 | F (10, 48) = 2.156 |
|  | *Garciellaceae* | P=0.4570 | F (10, 48) = 1.000 |
|  | *Lachnospiraceae* | ***P<0.0001*** | F (10, 48) = 7.101 |
|  | *Monoglobaceae* | P=0.0718 | F (10, 48) = 1.879 |
|  | *Butyricicoccaceae* | P=0.4430 | F (10, 48) = 1.018 |
|  | *Eubacterium coprostanoligenes group* | P=0.0049 | F (10, 48) = 3.018 |
|  | *Oscillospiraceae* | P<0.0001 | F (10, 48) = 8.065 |
|  | *Ruminococcaceae* | P=0.0056 | F (10, 48) = 2.963 |
|  | *UCG-010* | P=0.1842 | F (10, 48) = 1.459 |
|  | *Peptococcaceae* | P=0.0056 | F (10, 48) = 2.960 |
|  | *Anaerovoracaceae* | P=0.0015 | F (10, 48) = 3.533 |
|  | *Peptostreptococcaceae* | P=0.5594 | F (10, 48) = 0.8782 |
|  | *Peptostreptococcales-Tissierellales* | P=0.0700 | F (10, 48) = 1.890 |
| *Proteobacteria* | *Mitochondria* | P=0.5656 | F (10, 48) = 0.8711 |
|  | *Burkholderiaceae* | P<0.0001 | F (10, 48) = 25.64 |
|  | *Enterobacteriaceae* | P=0.3814 | F (10, 48) = 1.100 |
|  | *Xanthomonadaceae* | P=0.6153 | F (10, 48) = 0.8154 |
| *Verrucomicrobiota* | *Akkermansiaceae* | ***P=0.0087*** | F (10, 46) = 2.795 |

**Table S2: Summary of FluoroJade-C Staining.** Brain infection of male C57BL/6J mice with TMEV leads to an acute encephalitis phase and handling-induced symptomatic seizures. This acute seizure model induces acute hippocampal neurodegeneration that is detectable at 7 days post-infection, as assessed by FluoroJade-C histological staining (FJ-C) Data were compared within each brain region by proportion test, with * indicating significantly different from ABX-TMEV-CBZ treatment group, p<0.05, and ^#^indicating significantly different from SAL-TMEV-VEH treatment group, p<0.05

| **Treatment Condition** | | |  | **FJ-C Positive Labeling in:** | | | |
| --- | --- | --- | --- | --- | --- | --- | --- |
| **Gut Microbiome Depletion?** | **Infection** | **Treatment** | **Seizure History** | **CA1 (N/F)** | **CA3 (N/F)** | **DG (N/F)** | **Cortex (N/F)** |
| No (Saline) | Sham | CBZ | 0/5 | 0/5 | 0/5 | 0/5 | 0/5 |
| Yes (ABX) | Sham | CBZ | 0/5 | 0/5 | 0/5 | 0/5 | 0/5 |
| No (Saline) | TMEV | VEH | 7/7 | 7/7 | 7/7 | 0/7 | 5/7 |
| No (Saline) | TMEV | CBZ | 3/10^#^ | 4/10*^#^ | 6/10 | 0/10 | 4/10* |
| Yes (ABX) | TMEV | VEH | 5/7 | 7/7 | 7/7 | 0/7 | 7/7 |
| Yes (ABX) | TMEV | CBZ | 6/7 | 7/7 | 7/7 | 0/7 | 7/7 |

**Table S3: Liquid chromatography gradient for separation of carbamazepine in isolated mouse plasma following antibiotic-mediated gut microbiome dysbiosis in the Theiler’s virus mouse model of infection-induced acute seizures.**

| Time (min) | Flow Rate (mL/min) | Solvent A (%) | Solvent B (%) |
| --- | --- | --- | --- |
| Initial | 0.3 | 80 | 20 |
| 2 | 0.3 | 0 | 100 |
| 4.5 | 0.3 | 0 | 100 |
| 4.6 | 0.3 | 80 | 20 |

**Table S4: Mass spectrometry Transition for carbamazepine and diazepam-d5 (internal standard) quantification for isolated mouse plasma following antibiotic-mediated gut microbiome dysbiosis in the Theiler’s virus mouse model of infection-induced acute seizures.**

| **Compound** | **Parent (m/z)** | **Daughter (m/z)** | **Dwell time (s)** | **Cone (v)** | **Collision (eV)** |
| --- | --- | --- | --- | --- | --- |
| Carbamazepine | 237.0957 | 194.0025 | 0.096 | 2 | 18 |
| Carbamazepine | 237.0957 | 178.9191 | 0.096 | 2 | 32 |
| Diazepam-d5 | 290.1681 | 198.0652 | 0.096 | 100 | 36 |
| Diazepam-d5 | 290.1681 | 154.049 | 0.096 | 100 | 28 |

**Supplemental Figures.**

**Figure S1.** Hippocampal cell death, as assessed by FluoroJade-C staining (green) is seen in TMEV-infected mice, but not in sham-infected mice with and without a history of repeated antibiotics (ABX) administration. Blue indicates DAPI, a nuclear counterstain. Scale bar represents 100 μm.

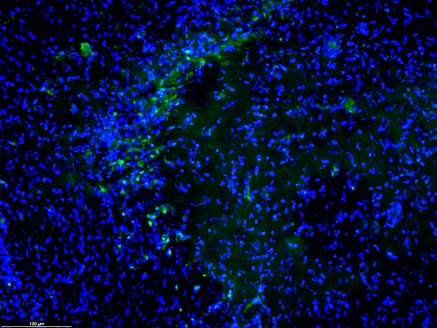

ABX, TMEV, CBZ

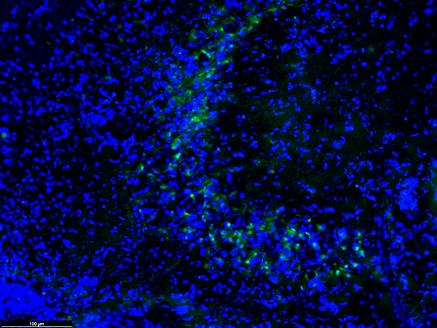

ABX, TMEV, VEH

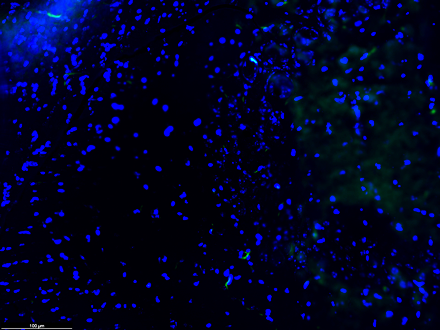

SAL, SHAM, VEH

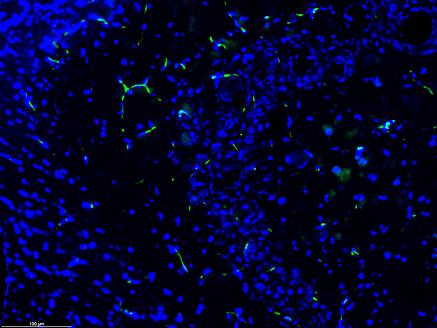

ABX, SHAM, VEH

**Supplemental Materials & Methods:**

Animal Handling and Diet Assignment. Male, wild type C57BL/6J mice (n=115; 4–5 weeks old; catalog #000664; Jackson Labs, Bar Harbor, ME) were group-housed throughout the infection and monitoring period (n=5/cage). Animals were given free access to autoclaved diet (ProLab RMH 3000) and filtered water except during periods of behavioral manipulation, as previously described^1^. Animals were maintained in standard housing chambers with corncob bedding in a temperature-controlled SPF vivarium on a 14:10 light/dark cycle (lights on: 6h00, lights off: 20h00) as previously detailed^2^. All studies were conducted between the hours of 9h00 and 17h00 during the animals’ light phase. This study was not designed to assess the impact of sex as a biological variable or the impact of sex hormones on seizure severity, thus only male mice were used. All animal use was approved by the University of Washington Institutional Animal Care and Use Committee (protocol 4387-02), conformed to the ARRIVE Guidelines, and was conducted in accordance with the United States Public Health Service's Policy on Humane Care and Use of Laboratory Animals.

Chemical and Reagents. *In vivo drug administration studies.* Methylcellulose (VEH; Sigma Aldrich catalog #M0430), carbamazepine (CBZ; Sigma Aldrich catalog #C4024). CBZ administered to mice was suspended in VEH. Ampicillin (Eugia US LLC, NDC 55150-114-20), Metronidazole (Azurity Pharmaceuticals, NDC 65628-0202), Neomycin Sulfate (Aspen Veterinary Resources Ltd., NDC 46066-211-07), and Vancomycin (Azurity Pharmaceuticals, NDC 65628-208-10) were dissolved in saline. *LC-MS/MS studies.* 7-chloro-1,3-dihydro-1-methyl-5-(phenyl-2,3,4,5,6-d_5_)-2H-1,4-benzodiazepin-2-one (diazepam-d5) was purchased from Cayman Chemical (Ann Arbor, MI). MS-grade acetonitrile (ACN) and methanol (MeOH) were purchased from Fisher Scientific (Pittsburgh, PA). Formic Acid (FA) 88% ACS grade was purchased from Sigma Aldrich (St. Louis, MO). All stock drug solutions, buffers, and HPLC mobile phase were prepared using Milli-Q grade water (Millipore, Bedford, MA). All other consumables were purchased from Fisher Scientific (Pittsburgh, PA). *Histopathology studies.* Fluoro-Jade C (FJ-C; Histochem catalog #2FJC), Shandon FormalFixx, (Fisher catalog #9990234), Potassium Permanganate (KMnO_4_; Sigma Aldrich catalog #223468), Acetic Acid (Sigma Aldrich catalog #695092), Hoechst 33342 nuclei acid stain (Invitrogen catalog #62249), phosphate buffered saline (PBS; VWR catalog #97062-948), DPX mounting media (Sigma Aldrich catalog #06522).

TMEV Infection*.* Mice were free-hand infected intracerebrally (i.c.) with either 20 mL Daniel’s (DA) TMEV strain, titer concentration of 3 x 10^5^ plaque-forming units (PFU) or sterile PBS under isoflurane anesthesia (Figure 1), as previously described^2^. TMEV titers were isolated from stock originally provided by Robert Fujinami at the University of Utah. All injection procedures were performed under sterile conditions. Following TMEV injection, the animals were monitored until they had recovered from anesthesia and were ambulatory.

Antibiotic Administration Prior to and During TMEV Infection: The oral ABX cocktail consisted of ampicillin (1 g/L), metronidazole (1 g/L), neomycin sulfate (1 g/L), and vancomycin (0.5 g/L) dissolved in saline^3^. Mice were pre-treated with ABX or saline (SAL) for 10 days beginning upon arrival at UW until conclusion of in-life testing 7 days p.i. (Figure 1). Mice were given 200 mL ABX or saline once daily, by oral gavage (Figure 1).

Carbamazepine Treatment During TMEV or SHAM Infection: Mice were administered CBZ (20 mg/kg) or VEH (0.5% methylcellulose) twice per day on days 3-7 p.i. (Figure 1). CBZ or VEH treatments were administered by the intraperitoneal (i.p.) route 30 minutes prior to each twice-daily behavioral seizure assessment (minimum 4 hours between sessions) during the acute TMEV infection period.

Assessment of Handling-Induced Seizures During TMEV Infection. TMEV and sham-infected mice were evaluated twice/day (minimum 4 hours between sessions), on days 3–7 p.i. for assessment of handling-evoked acute behavioral seizure severity (Figure 1). Seizure assessment involved brief cage agitation (<30 seconds), followed by individual handling of mice to visually assess righting reflex and gait. Typically, evoked secondarily generalized seizures occur in susceptible animals within this 5-10 minute observation period. The presence and severity of handling-induced seizures was scored according to the Racine scale (1- mouth and facial clonus; 2 – head bobbing; 3 – single forelimb clonus; 4 – bilateral forelimb clonus plus rearing; 5 – stage 4 plus repeated rearing and falling), as previously reported^4^ (Figure 1).

Complete Blood Counts at 7 Days Post-TMEV Infection: All TMEV- or sham-infected mice were euthanized via CO_2_ asphyxiation using a gradual displacement method (10%–30% volume per minute) according to the 2013 AVMA Guidelines for the Euthanasia of Animals (https://www.avma.org/KB/Policies/Documents/euthanasia.pdf). Blood was collected via cardiocentesis from a subset of mice immediately following euthanasia and placed directly into blood tubes containing K_2_EDTA (BD Microtainer Tubes with K_2_EDTA, catalog 365974; BD Biosciences, San Jose, CA) and serum separator tubes (BD Microtainer Serum Separator Tubes, catalog 365978; BD Biosciences), as previously described (Meeker et al, JPET 2019)^5^. Serum was separated by centrifugation (9000 relative centrifugal force x 5 minutes) within 2 hours of collection, and serum and whole blood from a subset of animals were submitted to a veterinary diagnostic laboratory for complete blood count and clinical serum biochemistry profiling (Moichor, San Francisco, CA).

Fecal Sampling Prior to and During TMEV Infection. Fecal samples were collected from a subset of animals in the study (n=5/experimental group). Time points of collection were: baseline/prior to housing within the University of Washington (UW) SPF vivarium; TMEV or sham infection day (0 p.i.), and conclusion of acute infection period (7 p.i.; Figure 1). Fecal sampling occurred by placing each mouse into an individual sterilized container, and spontaneous fecal samples were collected with a sterile needle. Fecal pellets were transferred into individual, sterile tubes, flash frozen, and stored at -80°C until processing. Baseline fecal sampling occurred prior to dietary or infection group randomization and thus reflected the baseline intestinal microbiome diversity upon arrival from Jackson Labs and prior to housing in the UW SPF vivarium. A total of 3 fecal pellets/mouse were obtained during the acute study period for bacterial 16S sequencing (Figure 1).

Fecal Sample 16S RNA Sequencing. Gut microbiome biodiversity was assessed by 16S rRNA amplicon sequencing of fecal samples. DNA was isolated from flash-frozen fecal pellets using a manual nucleic acid precipitation method at the University of Missouri Metagenomics Core Center (MUMC) as previously described^6^. Briefly, samples were placed in a 2 mL round-bottom tube containing 800 μL of lysis buffer and a sterile 0.5 cm diameter stainless steel bead and homogenized with a TissueLyser II.

Bacterial 16S rRNA amplicons were generated at the MUMC via amplification of the V4 hypervariable region of the 16S rRNA gene using single-indexed universal primers (U515F/806R) flanked by Illumina standard adapter sequences and the following parameters: 98°C(3:00) + [98°C(0:15)+50°C(0:30)+72°C(0:30)] × 25 cycles +72°C(7:00). Amplicons were then pooled for sequencing using the Illumina MiSeq platform and V2 chemistry with 2×250 bp paired-end reads, as previously described^6^. Samples returning greater than 10,000 reads were deemed to have successful amplification, consistent with established methods of the MUMC.

FluoroJade-C detection of cell death: Following in-life testing on Day 7 p.i., animals were euthanized by CO_2_ asphyxiation and brains of all mice were removed, flash frozen, and maintained at −80°C until histopathological processing. Two consecutive 20 μm-thick sections/mouse from the dorsal hippocampus (anteroposterior [AP] from Bregma: −1.34 mm) and ventral hippocampus (AP from Bregma: −2.24 mm) were sectioned on a cryostat (Leica DM1860) and slide-mounted. Slides were fixed in 4% Formal Fixx (10 min) and rinsed in DI water followed by FJ-C and Hoechst staining. Slides were imaged on an upright fluorescent microscope (Leica DM-4) with a 10x objective (40× final magnification) with constant acquisition settings, as previously conducted (Knox et al, Epilepsia 2021)^7^. Photomicrographs (n = 4 brain sections/mouse) were analyzed by an investigator who was blinded to treatment and visually scored for the presence of FJ-C positive cells in limbic structures (CA1, CA3, and dentate gyrus [DG] of dorsal hippocampus) and hypothalamic nucleus. If two or more of the consecutive sections from each mouse demonstrated FJ-C- positive staining in the designated brain region, the animal was counted as “FJ-C positive”.

Liquid Chromatography – Mass Spectrometry (LCMS) Quantitation of CBZ: Plasma was deproteinated prior to LCMS analysis. Subject plasma, calibration, or quality control samples (10 μL) were pipetted into a 1.2 mL Eppendorf microcentrifuge tube. Internal standard (10 μL) consisting of diazepam-d5 (1 mg/ mL in MeOH) was added to each microcentrifuge tube and vortexed for 5 seconds. Acetonitrile (ACN) (100 μL) was then added to each microcentrifuge tube and samples were vortexed for 15 seconds and then centrifuged at room temperature for 5 min at 16 RCF. Supernatant (50 μL) was removed and pipetted into 1 mL autosampler vials (Thermo Sciences, catalog #6PRV11-TR1 and 6PRC11STS1X). MilliQ water (450 mL) was added to each vial. Sample (10 mL) was injected onto the LC–MS platform. Chromatographic separation was achieved using a Water’s Acquity UPLC BEH C18 2.1x100 mm, 1.7 um (Waters Corporation, Milford, MA, USA) using a gradient consisting of 0.1% formic acid (FA) in H_2_O (A) and 0.1% FA in ACN (B) at a flow rate 0.3 mL/min (See Table S3). A Water’s Xevo-XS coupled to a Water’s I-Class Ultra high-pressure liquid chromatography system was used (Waters Corporation, Milford, MA, USA). Analytes were monitored in electrospray positive ionization mode (ESI+), using MRM mode. Instrument settings were as follows: Capillary (kV)—1.08, Source Temperature 150 °C, Desolvation Temperature 350 °C, Cone Gas Flow 150 L/hr, Desolvation Gas Flow 1000 L/hr, and Collision Gas Flow 0.16 mL/min. Two fragments for each analyte were used; one used for quantifying and the other as a qualifier. The m/z 237.1 > 194.0 and 178.9 transitions were used for carbamazepine. The m/z of 290.2 > 198.1 and 154.0 transitions were used for diazepam-d5 (see Table S4). Data was processed using MassLynx (Milford, MA) and linear equations were formulated by using peak area ratios (PAR’s) allowing for 15% variability across calibrators and quality controls for acceptability. Calibration Curve and QC samples were prepared by making dilutions of stock solutions containing carbamazepine from 10 mg/mL stocks in MeOH and stored at −20 °C. Calibration curves were created by analyzing drug-free plasma samples fortified with carbamazepine at 0.312, 0.625, 1.25, 2.5, 5, 10, and 20 mg/mL. Quality control (QC) samples in plasma were prepared at 1 and 5 mg/mL from separate dilutions of stocks than those used for the calibration curves.

Statistical Analysis: For in-life studies, average seizure burden and cumulative seizure burden were analyzed by repeat-measures two-way analysis of variance (ANOVA). Latency to first Racine stage 5 seizure was determined with a Kaplan-Meier plot. Spleen weights and complete blood counts were analyzed by two-way ANOVA. The number of animals with and without FJ-C positive labeling by histology within each brain region was analyzed by X^2^ test. Changes in microbiome commensal species were analyzed by two-way ANOVA. Changes in plasma CBZ concentrations were analyzed by two-way ANOVA. All statistical analysis was performed in GraphPad Prism version 10, with p < 0.05 considered significant.
